# Novel regional specific voltage-gated sodium channel (*vgsc*) mutations underlying pyrethroid resistance in *Aedes albopictus* (Skuse) from Northern Peninsular Malaysia

**DOI:** 10.1101/2025.03.17.643672

**Authors:** Nurul Adilah-Amrannudin, Kenneth Tan JunKai, Abu Hassan Ahmad, Hasber Salim, Mohd Ghows Mohd Azzam, Nurul Nadia Manap, Shamsudin Abdul Rahman, Intan Haslina Ishak

## Abstract

**Background:** Controlling *Aedes* mosquitoes remains as a primary means through the application of synthetic insecticides in Malaysia, but its efficacy is undermined by the evolution of resistance. *Aedes albopictus* as one of the dominant and competent arboviral vectors has an elusive insecticide resistance status in different geographical regions of Northern Peninsular Malaysia. This underscores the importance of assessing the diverse types of insecticides used and their association with target-site resistance mechanism in this species, which forms the basis of the present study.

**Methods:** WHO-bioassays were performed on *Ae. albopictus* larvae and adults from four localities (Penang and Perlis), towards 0.034 ppm temephos, 0.25% permethrin, 0.03% deltamethrin, 0.25% primiphos methyl and 0.1% propoxur. The partial *voltage-gated sodium channel* (*vgsc*) gene domain (DIIS6, DIIIS6, and DIVS6) of pyrethroid-exposed samples were subsequently genotyped through direct sequencing for any diagnostic single-nucleotide mutations, together with genetic variations and haplotype networks analysis. The predicted protein structures for the mutated regions and their binding affinities to pyrethroids were also evaluated using in-silico docking.

**Results and discussions:** Varying degrees of resistance were observed in all Penang and Perlis strains to all tested insecticides. Moreover, the detection of the F1534L mutation and newly discovered non-synonymous mutations (A1022S/P, E1041K, P1585R, and F1695L) suggest the progression of resistance alleles dissemination in other strains. The analysis of genetic variations, resistance allele distribution patterns, and haplotype networks showed evidence for multiple origins of these mutations. Data also revealed the discovered mutations affect the affinity of vgsc-binding proteins to pyrethroids.

**Conclusion:** This study highlights the genotype-phenotype associations in *Ae. albopictus* and their genetic links to pyrethroid resistance, offering insights to strengthen vector control strategies in Malaysia.

**Author summary:** In Malaysia, as in many other countries, dengue epidemic control primarily relies on managing the main vector, *Ae. albopictus* through commercially available insecticide applications. Vector control strategies have been extensively implemented by local health authorities, often without comprehensive information on insecticide resistance mechanisms in vector populations that could pose a major drawback of insecticide used. In an effort to assess the susceptibility status of *Ae. albopictus* populations, and determine the most effective insecticide for reducing Malaysian vector populations from different geographical settings, we conducted bioassays towards pyrethroid, organophosphate, and carbamate insecticide classes. Altogether, molecular-based assays were incorporated with phenotypic assay to elucidate the mechanisms modulating insecticide resistance and to unravel their genetic dynamics. Such screening offers detailed insights into specific mechanisms involved in conferring resistance for distinct adopted insecticides. This evidence aims to guide local health authorities in developing vector control strategies and exploring alternative solutions.

## Introduction

*Aedes albopictus* (Skuse), one of the most invasive mosquito species in tropical and subtropical regions, serves as a primary vector for dengue, chikungunya, and Zika virus [1]. *Aedes albopictus*, second only to *Ae. aegypti* in its role as dengue virus vector inflicts the disease onto millions of people in tropical and subtropical regions worldwide [2]. However, *Ae. albopictus* is gaining significance as a vector, with dengue virus detected in this arboreal mosquito, including in Malaysia [3], and despite a slight decrease in cumulative dengue cases from 120,418 in 2023 to 118,291 in 2024, there has been a continuing upward trend in related fatalities, rising from 93 to 111 deaths [4]. This species also possesses strong ecological adaptability, coupled with its ability to thrive near human dwellings, survive transport of its cold-hardy and drought-resistant eggs, and live in close association with humans, further underscores its importance [5], particularly in regard to its invasiveness and ability to displace *Ae. aegypti*.

Despite the introduction of Qdenga, the latest licensed dengue vaccine in Malaysia which is still in the process of gaining public confidence, vector control through the use of synthetic insecticide remains the goal standard method for controlling *Aedes* mosquitoes’ densities in the country [6–8]. In Malaysia, vector control programs often involve the use of pyrethroids, organophosphates, and propoxur, a practice carried out by the Ministry of Health, private pest control companies, and local communities [9,10] as well as in the agrarian sector [11]. These insecticides target the insect’s nervous system, albeit through different mechanisms, ultimately causing its death [12]. However, their prolonged and excessive use has fostered an overreliance on these insecticides, coupled with the improper application, which has contributed to the emergence of resistance in the mosquito vector [13].

The extensive use of insecticides is a major driver behind the growing reports of insecticide resistance in *Aedes* populations both in Malaysia and globally [14], jeopardizing the efficiency of existing vector control measures. In Malaysia, the primary vector, *Ae. aegypti*, has been the focus of numerous local studies in epidemic-prone cities [7, 15–18]. These studies have sparked interest in *Ae. albopictus* populations from various states, including Kuala Lumpur [8,16], Selangor [8,19], Perak [17], Sabah [20], as well as Pahang and Johor [8], highlighting the lack of comprehensive data on strains from Northern Peninsular Malaysia. This region, with its agricultural landscapes and diverse economic activities bordering Thailand, is prone to *Ae. albopictus* infestations in both urban and rural areas. Extensive insecticide use in these regions raises concerns about the emergence and evolution of insecticide resistance within this species.

Insecticide resistance primarily arises from two factors: changes in target-sites and an increase in the rate of insecticide metabolism [21]. Metabolic resistance is mainly driven by three enzyme families, namely cytochrome P450s, esterases, and glutathione S-transferases (GSTs), while target-site resistance is typically caused by one or more mutations in the insecticide’s target-site [13,22]. A key mutation linked to target-site resistance is the ‘knockdown resistance’ (*kdr*) mutation, which has been increasingly evolving in *Ae. albopictus*. This includes the emergence of V410L/A/G in Thailand and Italy [23], V1016G in China [24,25], India [26], and Europe [27], F1534S/C/L/W/I/R in Singapore [28], Indonesia [29] and China [23, 30–33], and I1532T in China [25,31,34], all of which are putatively confers resistance to DDT and pyrethroid insecticides.

Pyrethroid resistance is a major concern in Malaysia, where *Ae. albopictus* has been reported to have two identified mechanisms of resistance: target-site resistance (knockdown resistance, *kdr*) [19] and metabolic resistance [35]. The null detection of *kdr* mutations in *Ae. albopictus* from Malaysian in 2015 [16] was overturned five years later with the first discovery of the F1534L mutation in this species [19]. Even so, the genotypic screening of *kdr* mutations in Malaysian *Ae. albopictus* remains poorly defined despites various attempts at insecticide susceptibility profiling across different geographic regions of this species over time. Currently, there have been no attempts to investigate the distribution of alleles and genetic variations in the *vgsc* gene domains of *Ae. albopictus* from Malaysia that could influence the effectiveness of pyrethroid insecticides. Such fundamental insights are essential as an integral component of vector control programs and a prerequisite for effective interventions, forming the foundation of the present study on *Ae. albopictus* in Northern Peninsular Malaysia.

## Methods

### Study site

Field sample collections were carried out in residential areas in two states of Northern Peninsular Malaysia: Penang (Sungai (Sg.) Ara and Balik Pulau) and Perlis (Kampung Bahagia and Kampung Terus Teritip) (Fig 1). In each state, two localities with one classified as dengue outbreaks and the other as non-outbreak area were selected based on the iDengue Malaysia portal website (https://idengue.mysa.gov.my/) during the sample collection period from January 2022 till March 2023. A partnership was established with both Penang and Perlis Health District Officer to collaborate on site selection for mosquito sampling. The criteria for pre-selecting samples are as follows: residential areas with ongoing dengue cases for more than 14 days, extensive use of chemical insecticide control measures, and high *Aedes* density.

**Fig 1.**
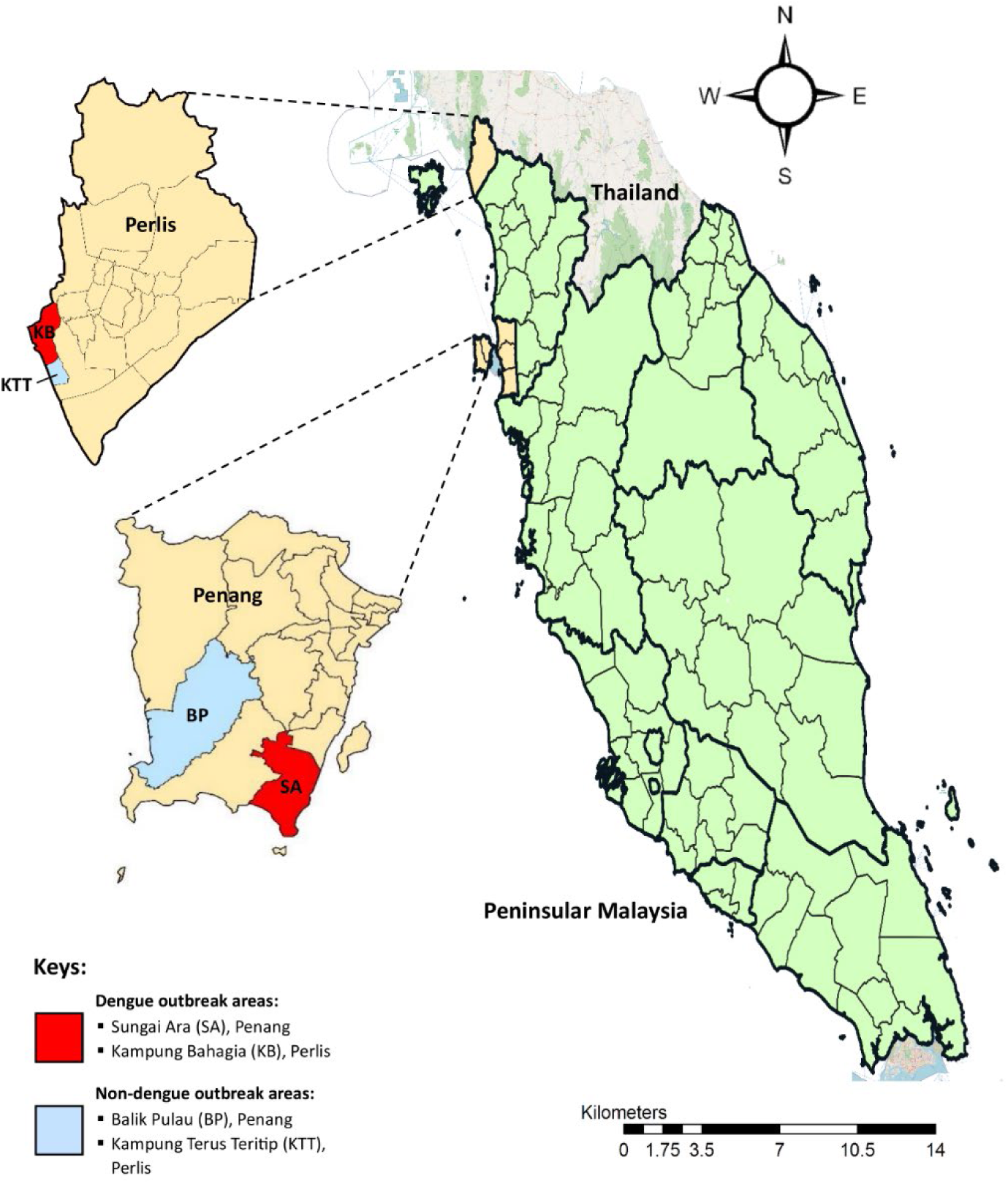
Geographical location of this study. The map of the study location in Northern Peninsular Malaysia (QGIS software).

### Mosquito collection

Sample collection was carried out using labelled ovitraps. An ovitrap is made up of a 400 ml black plastic container with a 6.5 cm aperture and base diameter and a 10 cm height. A rough-surfaced hardboard paddle (10 cm x 2.5 cm x 0.3 cm) was diagonally placed in each ovitrap to encourage mosquito egg-laying, and 5.5 cm of dechlorinated water was added to each trap. [36]. A total of 100 ovitraps were placed at random within and outside premises close to human habitation with the set-up distance of 5 to 10 m apart in each locality, in accordance with the regulation set forth by the Ministry of Health Malaysia [37]. After being deployed for 5-6 days, ovitraps were collected and transported back to the insectary. Additionally, *Aedes* larvae were also sampled from water-holding containers such as flowerpots, tree holes, and old tyres through pipetting and brought to the insectary for culturing purposes.

### Mosquito culture and establishment of F1 generations

All immature stages of *Aedes* mosquitoes were then reared to adults, and maintained in the standard conditions at 27±2℃ with the relative humidity of 75±10% and 12h:12h photoperiod [38]. They were fed with larval food containing ground cat biscuit, beef liver powder, milk powder and yeast with a ratio of 2:1:1:1. Emerging adults were morphologically identified to the species level using an appropriate key [39], and were fed with 10% sucrose solution ad libitum. Females aged four days were fed on a restrained rat (*Rattus norvegicus*) to generate F1 progenies. Three days post blood-feeding, a moist filter papers were folded into a conical shape and placed onto petri dishes filled to one-third with water as oviposit medium for the gravid females to lay eggs in 3-4 days, and the resulting F1 eggs were collected for experimentation. The susceptible laboratory strain obtained from the Vector Control Research Unit (VCRU) served as the reference strain.

### WHO susceptibility test

#### Larval bioassays

Larval bioassays were executed in accordance with WHO standard method [40] using F1 generation larvae. Bioassay was conducted in 250 ml of test solution with the appropriate concentration of the temephos were diluted in distilled water to produce the 6-8 discriminating concentrations ranging from 0.004 ppm to 0.042 ppm in a 16 oz plastic cup. For each concentration, 4 replicates of 25 healthy larvae were treated with temephos, alongside with two controls consisting of 1 ml of absolute ethanol in 249 ml distilled water. Larval mortality was recorded at 24-hour post-exposure. Larvae unable to swim to the surface were recorded as dead, while pupated larvae were excluded from the final count. The lethal concentration required to kill 50% and 95% of the tested samples (LC_50_ and LC_95_) was determined using probit analysis (IBM SPSS Statistics v27.0). Resistance ratios (RR) were calculated by comparing the LC_50_ and LC_95_ values to the *Ae. albopictus* USM VCRU susceptible strain. The standard procedures for larval bioassays were applied in the diagnostic dose experiments for both *Ae. albopictus* USM VCRU and field strains. The diagnostic dosage, set at twice the calculated LC_99.9_ of the USM VCRU susceptible strain, was used to assess the susceptibility of field strains.

#### Adult bioassays

The adult bioassays were conducted according to WHO protocol [40] using three –five-day-old non-blood fed F1 females *Ae. albopictus*. A total of 150 mosquitoes were exposed per insecticide, consisting of 4 replicates of 25 mosquitoes per tubes alongside two replicates for control. The insecticides that were tested are: 0.25% permethrin (Type I pyrethroid), 0.03% deltamethrin (Type II pyrethroid), 0.25% primiphos methyl (organophosphate), and 0.1% propoxur (carbamate), with replications of assay run for three times. The susceptibility bioassay test was conducted at 28 ± 2 °C with a relative humidity of 75 ± 10%. Post-treatment effect was observed after 24 hours, and the mortality was recorded. Both dead and alive mosquitoes were stored at -80°C for the molecular assays.

#### PBO-synergist bioassays

The effect of pre-exposure to the synergist, piperonyl butoxide (PBO) was also assessed for the population that showed mortality less than 98%. Female 3-5 days old non-blood fed F1 mosquitoes were exposed to papers impregnated with 4% PBO for one hour and subsequently exposed to pyrethroid insecticides: permethrin and deltamethrin using WHO susceptibility test. Mortality was recorded after 24 hours and compared to the results from each insecticide without PBO exposure, as well as to a control group exposed only to PBO.

### Genotyping of *kdr* mutations in *voltage-gated sodium channel* (*vgsc*) gene domains

#### Genomic DNA (gDNA) extraction

A total of ten gDNA of female *Ae. albopictus*, amounting five susceptible and five resistant mosquitoes from each locality, along with the USM VCRU susceptible strain for each pyrethroid insecticide, were individually extracted by adopting conventional extraction method based on Livak’s DNA Extraction protocol [41], with slight modifications. The DNA concentration and purity of extracted gDNA was measured using a Nanodrop spectrophotometer at 260 nm/280 nm and 260 nm/320 nm, and the samples were then stored at the -20°C for the downstream procedures.

#### Species identification

For each sample, the species identity of *Ae. albopictus* was confirmed using a PCR-based assay to amplify the ribosomal internal transcribed spacer *ITS1* with species-specific primers, as previously described by Ishak et al. [42].

#### *kdr* genotyping

Polymorphisms in the partial *vgsc* gene were genotyped in 90 females *Ae. albopictus* mosquitoes using three set of primers (S1 Table), representing each domain: domain II (DIIS6; exon 22 and 23), domain III (DIIIS6; exon 31), domain IV (DIVS6; exon 33). Primers were designed and adopted from Ahmad et al. [19] and Kasai et al. [28], and were synthesized commercially. The PCR mastermix was prepared using 2x Taq Plus Master Mix II (Vazyme, Nanjing, China). A total 25 μL of PCR mixtures comprised of 2 mM dNTPs, Vazyme Taq DNA polymerase, 6 mM MgCl2 (Vazyme Dye Plus), 10 μM of each primer set, and 100 ng DNA template. Thermocycling conditions for the PCR were 95°C for 3 minutes for initial denaturation, followed by 35 cycles at 95°C for 30 seconds, 55°C (DIIS6 and DIVS6) and 52°C (DIIIS6) for 30 seconds, 72°C for 1 minute and 1 cycle of final extension at 72°C for 5 minutes. The PCR analysis was run accordingly for the respective domain, and the amplified partial *vgsc* were subjected to electrophoretic separation and visualized on 1.5% (w/v) agarose gels pre-stained with RedSafe™ Nucleic Acid Staining Solution (INTRON Biotechnology, Korea). The PCR products were purified and directly sequenced in an automated Abi 3730xl Big Dye Terminator version 3.1 cycler (Applied Biosystems, Foster City, CA) using service from the Apical Scientific (Malaysia).

The chromatogram of each sequence was visually inspected using Chromas Lite v.2.1.1 (Technelysium Pty Ltd, Australia), and all 270 sequences (DIIS6, DIIIS6, and DIVS6) were compared with reference sequences deposited in the GenBank database to confirm their identity using Basic Local Alignment Search Tool (BLAST) (http://blast.ncbi.nlm.nih.gov.Blast.cgi). All sequences based on the respective domain were then assembled, aligned and trimmed using the ClustalW programs embedded in Molecular Evolutionary Genetics Analysis (MEGA) v11.0 software. Both synonymous and non-synonymous mutations, included *kdr* mutations were determined in this phase, and the sequences were deposited in the NCBI GenBank under accession number PQ660263-PQ660378.

#### Statistical analysis

The percentage mortality 24 hours post-exposure was used to determine the phenotypic status of each population. Susceptible and resistant phenotypes were classified based on WHO guidelines. A strain was classified as susceptible if the mortality rate was ≥ 98%, as possibly resistant if the mortality rate ranged between 90% and 97%, and as resistant if the mortality rate was < 90%. Odds ratio (OR) analysis and Fisher’s exact test were performed to compare the distribution of *kdr* or other genotypes in *vgsc* gene of both dead and alive mosquitoes. Comparative measure of knockdown and mortality between all *Ae. albopictus* strains in this study were conducted using one-way analysis of variance (ANOVA) [43,44]. Cross-resistance among insecticides was investigated by analyzing the associations between the mortality rates of tested adulticides using Pearson correlation analysis, with P-values ≤ 0.05 considered statistically significant. All statistical analysis were performed using IBM SPSS Statistics v27.0.

### Genetic diversity analysis of *vgsc* gene domains

DNA polymorphism in the *vgsc* gene domains, based on interpopulation variability, was assessed in terms of the number of polymorphic sites, parsimony-informative sites, haplotype diversity (Hd), nucleotide diversity (π), the number of segregating sites (S), the average number of nucleotide differences (K), and neutrality tests (Tajima’s D) across the alignment of all 270 sequences (DIIS6, DIIIS6, and DIVS6). These analyses were conducted according to geographical origins using the DnaSP v.5.10.1 software [45].

### Haplotype networks

A haplotype network based on the partial *vgsc* gene domains was constructed using Population Analysis with Reticulate Trees (PopART) (http://popart.otago.ac.nz/index.shtml) to assess the genealogical relationship of *Ae. albopictus vgsc*-kdr haplotypes. Haplotypes were linked from the shortest to the longest distance until all were integrated completely. Population divergence for probabilities were analyzed using TCS networks inference [46] constructed within the PopART software.

### Homology modelling of partial *vgsc* gene domain’s protein

The 3D protein structure of partial *vgsc* DIIS6, DIIIS6 and DIVS6 proteins were constructed by ColabFold v1.5.5: AlphaFold2 using MMseqs2 [47] pipeline and default AlphaFold parameters. Five predicted models generated for each model will then be subjected to PROCHECK in UCLA-DOE LAB — SAVES v6.1 (https://saves.mbi.ucla.edu/) for Ramachandran plot analysis for structural validation. Models were assessed with the predicted local distance difference test (pLDDT) scores >70 from AlphaFold that indicate high confidence for domain-specific structures models, and relatively better Ramachandran plot statistics with the highest percentage of residues in allowed regions were prioritized (S1 Fig). Models with fewer residues in disallowed regions and relatively higher pLDDT scores was favoured.

### Docking

3D conformers of deltamethrin and permethrin were obtained from PubChem compound (https://pubchem.ncbi.nlm.nih.gov/docs/compounds), and were set as ligands following the energy-minimization using the MMFF94 force field in Open Babel. Following the selection of optimal protein model, docking simulation was performed with AutoDock Vina integrated within PyRx 0.8 [48]. Binding affinities (in kcal/mol) and docked poses with the lowest RMSD values were recorded. To assess binding affinity differences between mutant proteins, the best docked post for wild-type and mutant proteins were aligned and visualized in UCSF ChimeraX [49].

## Results

### Susceptibility profiling to insecticides

#### Susceptibility status of larval *Ae. albopictus*

*Aedes albopictus* larvae were subjected to 0.012 ppm and 0.034 ppm temephos larvicides, showing very low mortality (ranging from 5.67% to 59.00%) across all field strains (S2 Table; S2 Fig). Slightly higher of RR_50_ and RR_95_ were observed in *Ae. albopictus* from KB strains from Perlis, as compared to other strains (S3 Table).

#### Susceptibility status of adults *Ae. albopictus*

##### Mortality rates and cross-resistance correlation

All the field strains of *Ae. albopictus* from Penang and Perlis showed a varying degree of resistance against carbamate, organophosphate and pyrethroids (S4 Table). Percentage mortality varied from 2.00% to 19.33% for propoxur, 70.00% to 94.33% for primiphos methyl, 34.00% to 93.00% for permethrin, and 3.67% to 96.67% for deltamethrin, accordingly. All insecticides were indiscriminately lethal towards susceptible USM VCRU populations, resulting in complete mortalities overall (Fig 2). The lowest percentage of mortality toward propoxur was observed in Kampung Bahagia (KB), Perlis, with 2.00% mortality. Resistance to primiphos methyl was shown in all populations, with the highest lethal rates in Kampung Terus Teritip (KTT) strain in Perlis (70.00%) and in Balik Pulau (BP), Penang (92.33%). In pyrethroid bioassay, *Ae. albopictus* from Penang exhibited the intensity of low to moderate resistance towards permethrin (Sungai Ara (SA): 79.00%; BP: 93.00%) and deltamethrin (SA: 96.67%; BP: 90.00%). As for Perlis strains, dissimilitude in mortalities were observed since they are more formidable than Penang strains when being exposed to permethrin (SA: 34.00%; BP: 48.67%) and deltamethrin (SA: 3.67%; BP: 10.50%). Pearson’s correlation analysis revealed a significant correlation between permethrin and deltamethrin usage (r = 0.950, P = 0.050), indicating cross-resistance. However, no meaningful correlation was found between temephos (0.034 ppm) and primiphos-methyl exposures (r = -0.112, P = 0.888). Moderate inter-class cross-resistance was observed between primiphos-methyl and propoxur (r = 0.607, P = 0.393).

**Fig 2.**
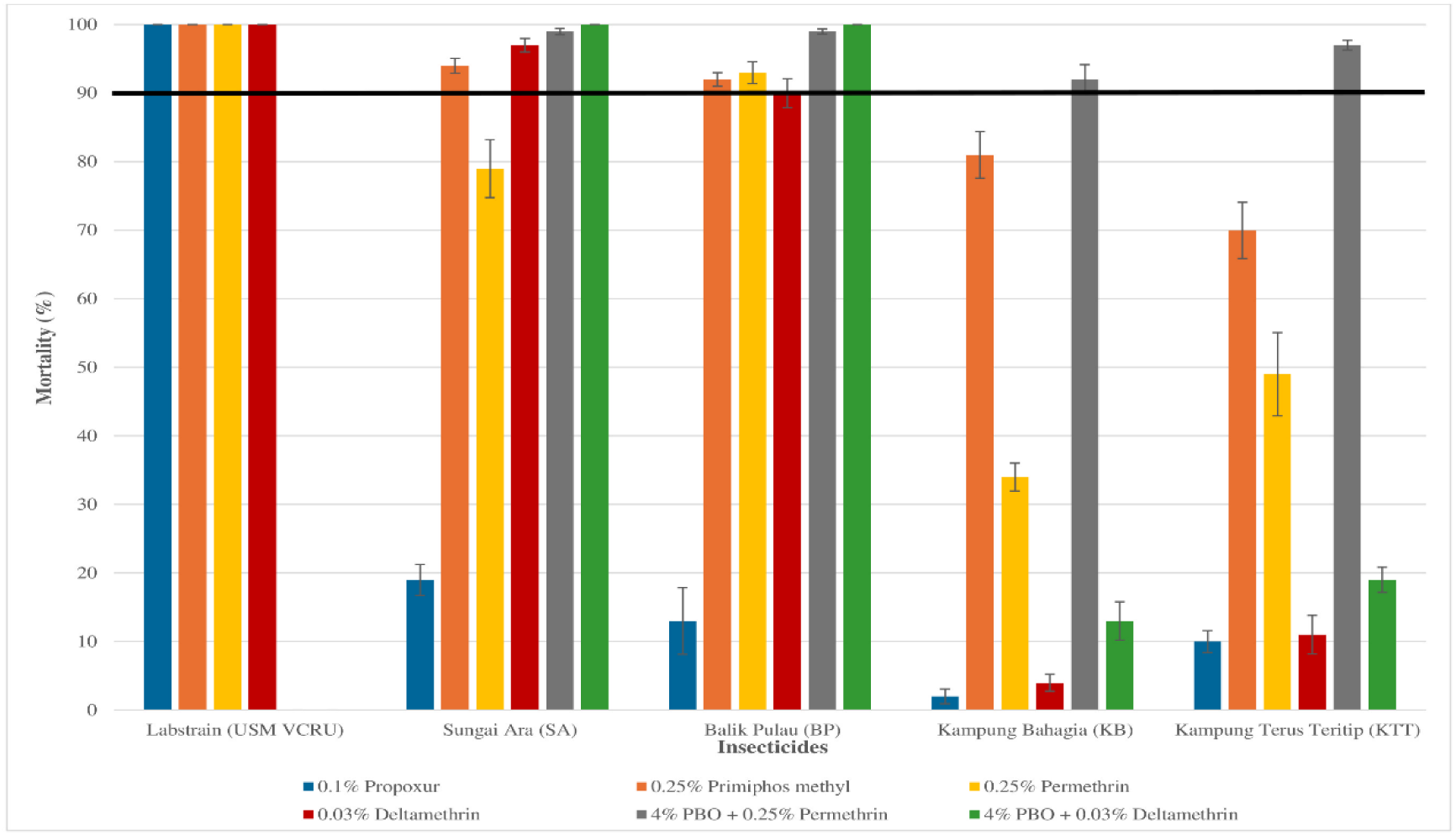
Percentage mortality of female *Ae. albopictus* from four different localities in Penang and Perlis, Malaysia exposed to three classes of insecticides and a PBO-synergist. Error bars represent 95% confidence interval (CI). Black horizontal line indicates resistant threshold level (mortality <90% is considered phenotypically moderate to high resistant-*Ae. albopictus*; WHO, 2016).

##### Knockdown effects

The knockdown times for 50% (KdT_50_) and 95% (KdT_95_) of *Ae. albopictus* were calculated for pyrethroid exposure. Heterogeneity in knockdown rates of the field strains *Ae. albopictus* can be observed with various degrees of resistance folds, with varying knockdown rates at KdT_50_ and KdT_95_ (KdT_50_: 15.57 to 199.45 minutes; KdT_95_: 28.22 to 378.38 minutes) (S5 Table). Deltamethrin was the most effective insecticide, with the shortest KdT_50_ and KdT_95_ times, knocking down 50% of *Ae. albopictus* in 15.57 minutes (SA) and 16.86 minutes (BP), while permethrin was the least effective in Penang populations. Conversely, deltamethrin depicted very low efficacy towards Perlis’ *Ae. albopictus* populations as this insecticide incapable at inducing high knockdown percentage (>5%), rendering to inability to generate KdT_50_ and KdT_95_ values, indicating high resistance. In different circumstances, knockdown rates were yielded from permethrin usage-imposed Perlis populations.

##### Synergism assay with PBO

Pre-exposure to 4% PBO for 1 hour prior to permethrin exposure increased mortality rates across all populations. All Penang strains showed a recovery of 99.00% in SA and 99.50% in BP after exposure to PBO and permethrin, with full recovery observed in both strains when pre-treated with PBO and deltamethrin. Pertaining to Perlis strains, profound synergistic effect was obtained after exposure to PBO and Permethrin, amounting mortalities from 34.00 to 91.50%, and 48.67% to 97.00% for KB and KTT, respectively. On the contrary, very low restoration of susceptibility was shown amongst both strains (KB: 3.67% to 12.50%; KTT: 10.50% to 19.00%) at post-exposure of PBO and deltamethrin. The changes in the percentage mortality of mosquito samples that were pre-treated with a PBO-synergist following pyrethroids is shown in both S2 Table and Fig 2. Pearson’s analysis showed a positive but non-significant correlation between permethrin-treated *Ae. albopictus* mortality rates, with and without PBO (r = 0.898, *P* = 0.102; *P* ≤ 0.05), while Spearman’s test revealed a strong association for deltamethrin-treated mosquitoes pre-exposed to PBO (r = 0.949, *P* = 0.051; *P* ≤ 0.05). Different correlation coefficient methods were used due to the data in the latter test not normally distributed.

### Genotyping of *kdr* mutation of *Ae. albopictus* across Northern Peninsular Malaysia

#### Detection of *kdr* mutation in the *vgsc* gene domains of *Ae. albopictus*

Ninety *Ae. albopictus* samples were randomly selected from pyrethroid Type I and II resistant and susceptible populations from the field strains to determine the presence of *kdr* mutations at codons 989, 1011, 1016, 1520, 1532, 1534, and 1763 in DIIS6, DIIIS6, and DIVS6, together with their allelic frequencies. This finding revealed the presence of only the F1534L mutation, characterized by allelic changes from TTC/TTC (wild-type; phenylalanine) to TTC/TTG (heterozygous mutant; leucine), in both the Penang and Perlis populations. The heterozygous mutation was notably more prevalent in Perlis strains (32.5%) compared to Penang strains (5%). This *kdr* mutation was predominantly identified in KB-resistant and susceptible *Ae. albopictus*, with 35% and 15% L/TTG detection rates and allelic frequencies of 0.6 and 0.4, respectively. Whilst in Penang, it was observed at a prevalence of 10%, exclusively in the resistant SA strain.

Multiple novel non-synonymous mutations were detected in all studied domain, mostly were discovered in resistant *Ae. albopictus*. In DIIS6, amino acid substitution of alanine (GCC, wild type) to serine (TCC) and proline (CCC) at codon 1022 was circulated within populations from Penang and Perlis. The heterozygous A1022S mutation was frequently identified in Penang strains (17.5%), whereas the homozygous A1022S mutation was only found in Perlis strains (10%). The highest distribution was observed in the SA population, with frequencies of 25% in resistant and 5% in susceptible *Ae. albopictus*, respectively. Interestingly, A1022P genotype was only circulated in Penang populations, where this homozygous variant was found in resistant mosquitoes from BP (20%) and SA (10%). Furthermore, a substitution of glutamic acid (GAA, wild type) to lysine (AAA) at codon 1044 was solely found in resistant *Ae. albopictus* from Penang and Perlis, with the highest homozygous mutation was detected in SA strain (40%), contributing to an allelic frequency of 0.6 in deltamethrin-treated mosquitoes.

In DIIIS6, for new mutant genotype P1585R, denoting amino acid alteration of proline to arginine (CCG to CGG, or CGC). This homozygous R/CGG variant was detected with a prevalence of 50% in resistant-SA strains from Penang and 60% across both resistant and susceptible KB and KTT strains, with 20% occurrence of R/CGC observed in Perlis strains overall. Meanwhile, in DIVS6, there is no identification of *kdr* 1763 alleles, but there is a novel occurrence of L/CTC allele across all resistant *Ae. albopictus* though, which was quite high in SA (50%) and BP (30%). The occurrences of all related alleles referring the genotypic prevalence in DIIS6, DIIIS6, and DIVS6 of the studied *Ae. albopictus* were illustrated in Fig 3, along with their allele frequency rates shown in Fig 4. Altogether, the occurrence of mutant alleles in *Ae. albopictus* strains from Northern Peninsular Malaysia made up 15.5% of the total detections (S6 Table), and percentile of genotypic distributions in both Penang and Perlis populations were summarized in Fig 5.

**Fig 3.**
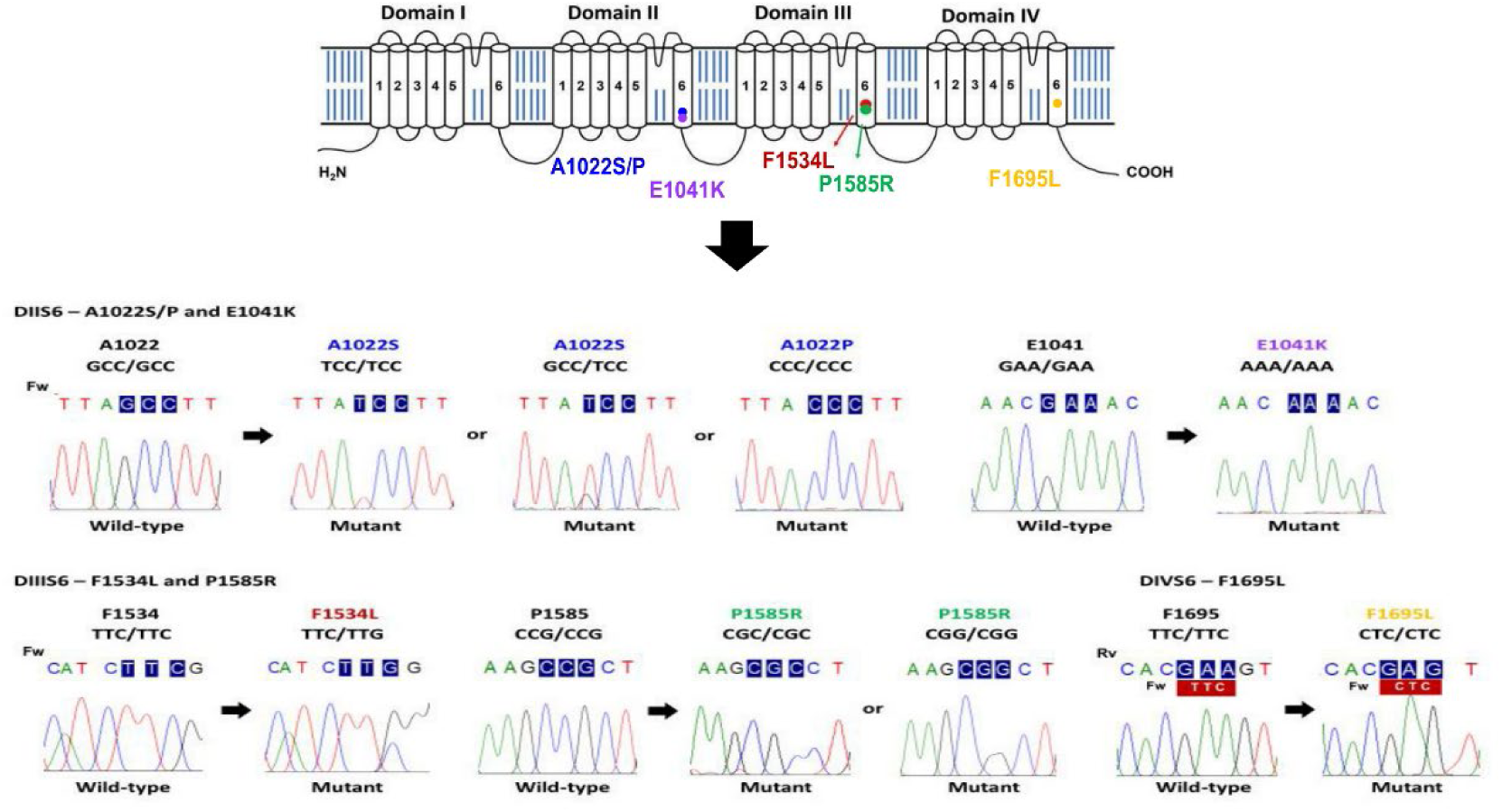
Chromatogram of the wild-type and mutated alleles found in this study. Schematics diagram of voltage-gated sodium channel (*vgsc*) with the occurrence of several mutations in the studied *Ae. albopictus* illustrated by chromatogram of sequence data, displaying both wild-type and mutated alleles that could affect the binding of pyrethroid insecticides to the *vgsc* target-site.

**Fig 4.**
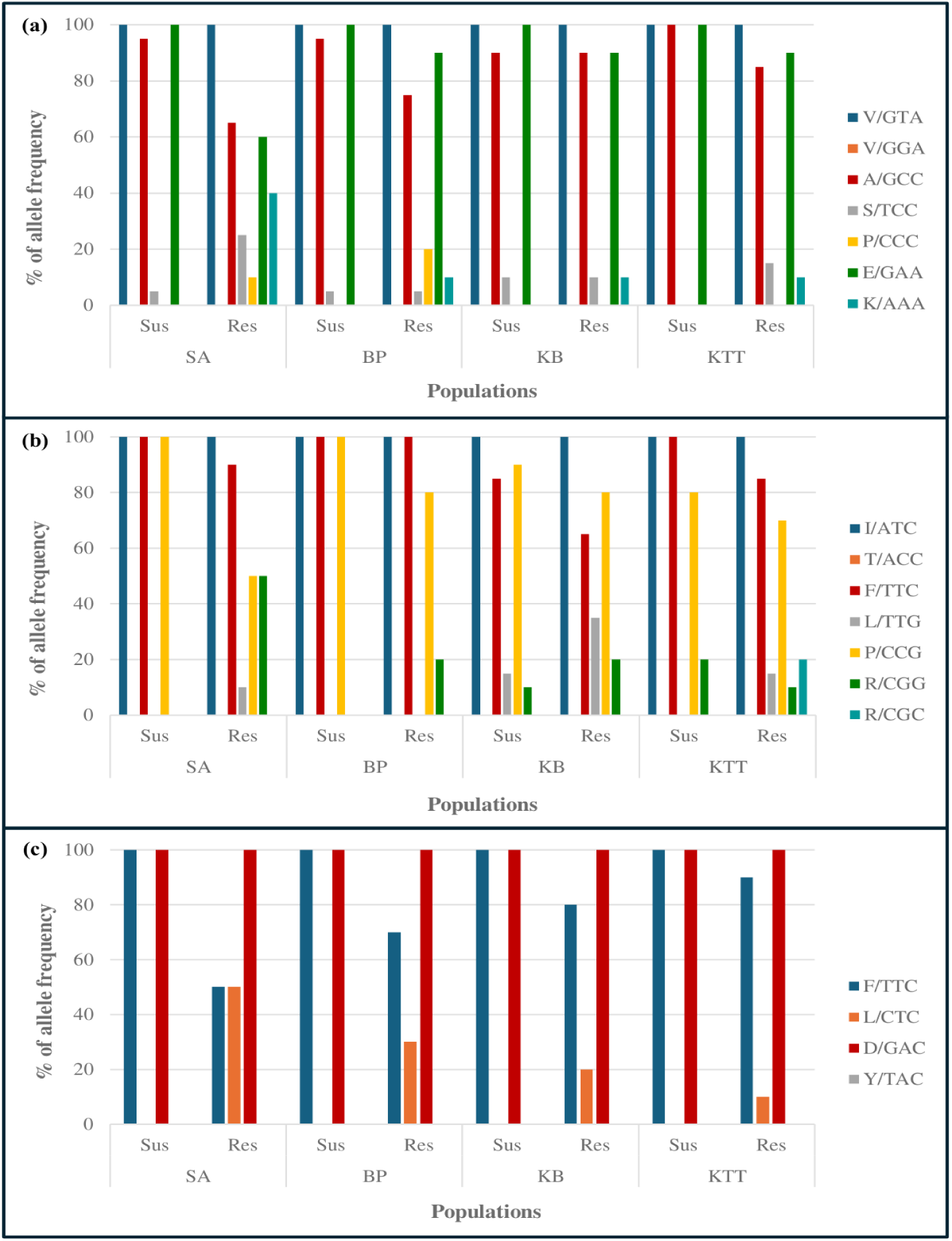
Percentage of allele frequency. Percentage of allele frequency at (a) Domain IIS6 comprised of codons 1061, 1022, and 1041, at (b) Domain IIIS6 comprised of codon 1532, 1534, and 1585, and at (c) Domain IVS6 comprised of codon 1695 and 1763 of the voltage-gated sodium channel (vgsc) gene in *Ae. albopictus* populations.

**Fig 5.**
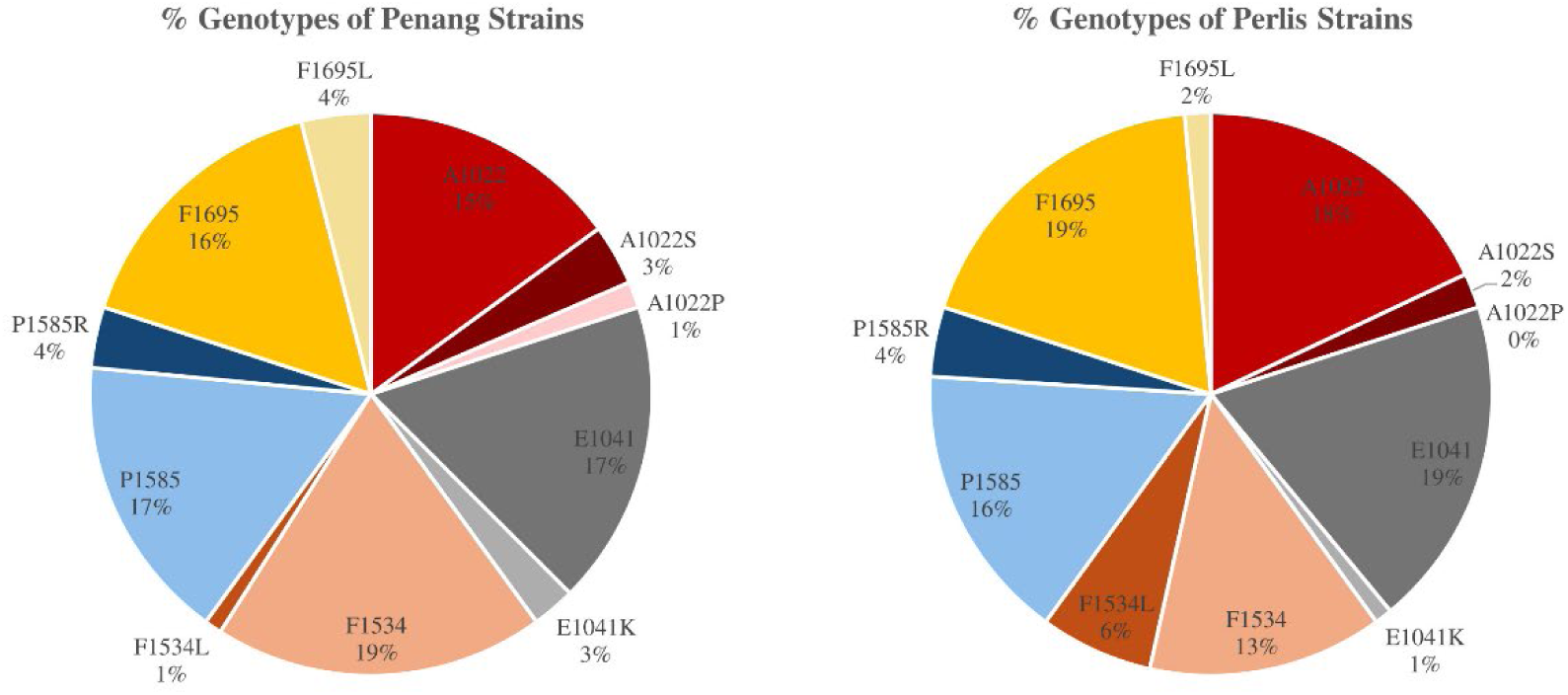
Genotypic distributions. The percentile of genotypic distributions associated with susceptibility profiles of *Ae. albopictus* in both Penang and Perlis populations.

### Correlation between putative *kdr-*1534 and newly discovered non-synonymous mutations at DIIS6, DIIIS6, and DIVS6 with pyrethroid resistance

The impact of putative *kdr* mutations (F1534L) and newly discovered non-synonymous mutations (A1022S/P, E1041K, P1585R, and F1695L) on pyrethroid resistance was assessed by examining the association between these mutations and both the permethrin and deltamethrin resistant phenotype through genotyping 90 resistant and susceptible *Ae. albopictus* across all populations at DIIS6, DIIIS6, and DIVS6. For the putative kdr-1534 variant, allele frequencies ranged from 0.0 to 0.4 across all studied populations. SA population showed the potential of heterozygote F/L to cause permethrin resistance (OR = 6.177, P-value = 0.0474; P<0.05), whilst the absence of this genotype does not impact resistance amongst BP populations towards both insecticides as well as SA to deltamethrin (OR = 1.000, P-value = 1.000; P<0.05). On the contrary, there was the occurrence of F1534L mutation in both susceptible and resistant *Ae. albopictus* strains in Perlis with OR varies from 2.667 to 6.177 (S7 Table). This implies the individuals with the mutation are more likely to be resistant, although not statistically significant that probably owing to the small sample size.

Meanwhile, for new mutations at codon 1022 in DIIS6, the resistant *Ae. albopictus* depicted a high likelihood of pyrethroid resistance, with a marginally increased frequency of P/P homozygote resistant genotype, and its counterparts of the heterozygote genotype A/S. The OR ranged from 3.857 to 14.539, despites a weak association with the resistant phenotype (P<0.05) (S8 Table). This finding was in align with the E1041K mutation that positively linked to deltamethrin resistance in Penang (only SA) and Perlis strains, albeit less significant. However, a higher frequency of K/K homozygote resistant genotype confirmed a strong correlation between E1014K genotypes and deltamethrin resistance, particularly in the SA population of Penang (OR = 30.333, P = 0.011; P<0.05) (S9 Table). For P1585R, the positive correlation of high mutant alleles frequencies, insuring the strong correlation between R/R homozygote genotype with deltamethrin resistant individuals in SA population, Penang (OR = 30.333, P-value = 0.011; P<0.05) (S10 Table). The presence of this genotype was more likely to exhibit resistance likelihood towards both insecticides among individuals in Penang as opposed to the Perlis strains, in which this genotype was less affected the permethrin-treated individuals (OR = 0.162 to 1.000). Whilst, in F1695L genotyping, the presence of this genotype statistically shows the susceptibility of SA strain towards permethrin predominantly (OR = 71.400, P-value = 0.011; P<0.05). Based on the S11 Table, this homozygote genotype portrayed high tendency in causing resistance towards permethrin compared to deltamethrin with varying OR values from 14.539 to 71.400. In Perlis populations, all strains showed resistance likelihood to permethrin and deltamethrin, except for KTT, where the F1695L variant showed no correlation with susceptibility or resistance (OR = 1.000).

### Distribution of single-, duo-, triple-, and quadruple-loci of the genotypic combination in DIIS6, DIIIS6, and DIVS6 of *vgsc* gene in *Ae. albopictus* and its association with pyrethroid resistance

Amongst the 80 field-strain *Ae. albopictus* from Northern Peninsular Malaysia genotyped for DIIS6, DIIIS6, and DIVS6, 15 distinct substitution patterns were identified, including 5 single locus (24.44%), 6 duo-locus (10%), 2 triple-locus (2.22%), and 3 quadruple-locus (3.33%) combinations (S12 Table). The occurrence of single mutations of A1022S/P, E1041K, F1534L, P1585R, and F1695L dominantly prevailed with the frequency rates of 17.78% in resistant and 6.67% in susceptible mosquitoes. Single mutations occurred in 14.44% of Perlis strains and 10% of Penang strains, whereby F1534L was exhibited in 7 resistant samples from Perlis. Meanwhile, duo-loci mutations were observed with 5 unique patterns that involved the combinations of amino acid changes in both similar and different partial gene domains of all resistant samples, as tabulated. We noticed that E1041K variant consistently co-existed with other mutations, suggesting its potential role in these co-occurrences. The triple mutant combination of A1022P + E1041K + P1585R were only seen in two SA-resistant deltamethrin-treated samples. Furthermore, four combinations of quadruple-loci genotypes were particularly present with two distinct combinations: A1022P + E1041K + P1585R + F1695L, detected in SA permethrin-resistant sample, and A1022S + E1041K + P1585R + F1695L, identified in deltamethrin-resistant samples of KB and KTT, respectively.

Overall OR analysis of all combinations of genotypes in *Ae. albopictus* from Northern Peninsular Malaysia depicted strong associations of these genotypes to pyrethroid resistance (Table 1). *Aedes albopictus* from Penang populations demonstrated significantly stronger associations of allele changes in conferring resistance compared to Perlis strains (Penang: OR = 36.616, P = 0.0001; Perlis: OR = 3.434, P = 0.0153; P < 0.05). It can be inferred from the presence of higher frequencies of homozygote genotypes in Penang populations, contradicting to the Perlis strains which mostly affected from the occurrences of heterozygote genotypes mutations in DIIIS6. To sum up, a cumulative genotyping of all mutations discovered in the studied *Ae. albopictus* that represented Northern Peninsular Malaysia strains (OR = 9.120, P-value = 0.0001; P<0.05), were strongly associated to both the resistant phenotypes and also affects the susceptible individuals.

**Table 1.** Summary of all prevalent genotypes related to *vgsc* mutations with the resistance phenotype towards pyrethroid insecticides for the Penang and Perlis strains of *Ae. albopictus*.

| No of alleles |  |  |  |  |  |  |  |  |  |  |  |  |  |  |  |  |  |  |
| --- | --- | --- | --- | --- | --- | --- | --- | --- | --- | --- | --- | --- | --- | --- | --- | --- | --- | --- |
| Strain | Phe | n | DII |  |  |  |  | DIII |  |  |  |  | DIV |  | Mutant allele frequency | % of mutant allele (%) | Odd-ratio (OR) | Fisher's Exact test (p-value) |
|  |  |  | A1022S/P |  |  | E1041K |  | F1534L |  | P1585R |  |  | F1695L |  |  |  |  |  |
|  |  |  | <u>G</u>CC | <u>T</u>CC | <u>C</u>CC | <u>G</u>AA | <u>A</u>AA | <u>T</u>TC | <u>T</u>TG | <u>C</u>CG | <u>C</u>GG | <u>C</u>GC | <u>T</u>TC | <u>C</u>TC |  |  |  |  |
| SA | Sus | 10 | 19 | 1 | 0 | 20 | 0 | 20 | 0 | 20 | 0 | 0 | 20 | 0 | 0.1 | 19 | 58.143 | 0.0001** |
|  | Res | 10 | 13 | 5 | 2 | 12 | 8 | 18 | 2 | 10 | 10 | 0 | 10 | 10 | 3.7 |  |  |  |
| BP | Sus | 10 | 19 | 1 | 0 | 20 | 0 | 20 | 0 | 20 | 0 | 0 | 20 | 0 | 0.1 | 9 | 20.277 | 0.0001** |
|  | Res | 10 | 15 | 1 | 4 | 18 | 2 | 20 | 0 | 16 | 4 | 0 | 14 | 6 | 1.7 |  |  |  |
| Penang | Sus | 20 | 38 | 2 | 0 | 40 | 0 | 40 | 0 | 40 | 0 | 0 | 40 | 0 | 0.1 | 14 | 36.616 | 0.0001** |
|  | Res | 20 | 28 | 6 | 6 | 30 | 10 | 38 | 2 | 26 | 14 | 0 | 24 | 16 | 2.7 |  |  |  |
| KB | Sus | 10 | 18 | 2 | 0 | 20 | 0 | 17 | 3 | 18 | 2 | 0 | 20 | 0 | 0.7 | 13 | 3.116 | 0.0193* |
|  | Res | 10 | 18 | 2 | 0 | 18 | 2 | 13 | 7 | 16 | 4 | 0 | 16 | 4 | 1.9 |  |  |  |
| KTT | Sus | 10 | 20 | 0 | 0 | 20 | 0 | 20 | 0 | 16 | 4 | 0 | 20 | 0 | 0.4 | 10 | 4.571 | 0.0081* |
|  | Res | 10 | 17 | 3 | 0 | 18 | 2 | 17 | 3 | 14 | 2 | 4 | 18 | 2 | 1.6 |  |  |  |
| Perlis | Sus | 20 | 38 | 2 | 0 | 40 | 0 | 37 | 3 | 34 | 6 | 0 | 40 | 0 | 0.6 | 12 | 3.434 | 0.0153* |
|  | Res | 20 | 35 | 5 | 0 | 36 | 4 | 30 | 10 | 30 | 6 | 4 | 34 | 6 | 1.8 |  |  |  |
| NPM | Sus | 40 | 76 | 4 | 0 | 80 | 0 | 77 | 3 | 74 | 6 | 0 | 80 | 0 | 0.3 | 27.5 | 9.120 | 0.0001** |
| strains | Res | 40 | 63 | 11 | 6 | 66 | 14 | 68 | 12 | 56 | 20 | 4 | 58 | 22 | 2.2 |  |  |  |

Phe.: Phenotype; Range of odd ratio (OR): OR = 1 (No association between genotype and phenotype), OR <1 (Negative association, or low likelihood of insecticide resistance), OR >1 (Positive association, or high likelihood of insecticide resistance). \*Statistical significance (Fisher's Exact test): P<0.05; \*\*: extremely significant, \*: significant.

### Haplotype distributions and polymorphism analysis of the partial *vgsc* gene domains (DIIS6, DIIIS6, and DIVS6) in Northern Peninsular Malaysian of *Ae. albopictus*

#### *kdr-vgsc* haplotypes

The genealogical network identified 41 haplotypes, 25 haplotypes, and 50 haplotypes based on the partial analysis of 326 bp, 221 bp, and 283 bp of DIIS6, DIIIS6, and DIVS6 of 90 *Ae. albopictus* samples, respectively (S13 Table). All of these haplotypes were deposited in the NCBI GenBank. In DIIS6, 45 polymorphic sites were identified, leading to the formation of 41 haplotypes of wild-type putative *kdr* alleles. H3 is the largest and most connected ancestral haplotypes (16.67%), comprising of the wild-type alleles with genotype V1016, A1022, and E1041 dominated with USM VCRU susceptible strain, suggesting that these haplotypes were prevalent before the onset of insecticide selection pressures. Whilst a total of 7 haplotypes consisted of mutant genotype A1022S/P and 6 haplotypes carried both genotype A1022S/P and E1041K (S2 Fig), implying the distributions of these predictive-resistance alleles in *Ae. albopictus* colonized in both Penang and Perlis. For DIIIS6, *kdr* mutations at codon 1534 were distributed in different haplotypes, signifying independent origins (Fig 6).

**Fig 6.**
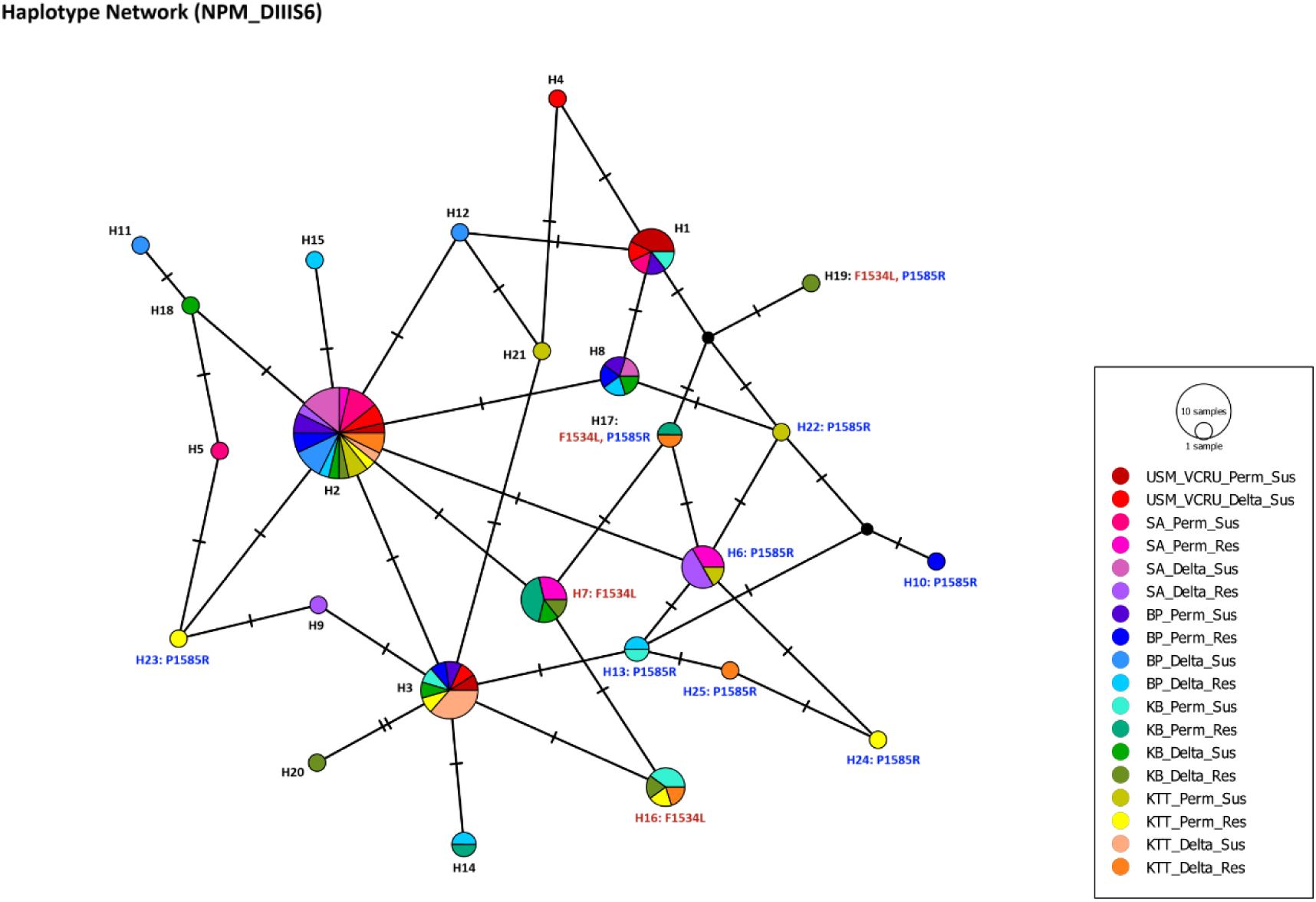
Haplotype network. Haplotype network showing the genealogical relationships of 25 haplotypes in DIIIS6 based on four strains of Northern Peninsular Malaysia populations of *Ae. albopictus*: wild-type haplotypes are depicted in black, haplotypes with *kdr* mutations associated with resistance phenotypes are marked in red, and those with new non-synonymous mutations are highlighted in blue.

The haplotypes carrying *kdr* allele (H7 and H16) and P1585R predictive-resistance allele (H17 and H19) were observed to be connected through a series of mutational events. kdr-1534L haplotype, H7 was derived from the ancestral haplotypes H2 (31.11%) subsequently diverged into H16, H17, and eventually H19. Interestingly, the divergent haplotypes were exclusively comprised of Perlis samples. Besides that, most of the unique haplotypes (H1, H5, H9, H10, H15, H19, H20, H23, H24, H25), which evolved from their distinct ancestors, represent resistant mosquitoes. This finding could imply the uniqueness of resistant trait towards a certain selection pressure. The closely related haplotypes of DIIIS6 inferred recent evolutionary events of *Ae. albopictus*. Although no D1763 *kdr* haplotypes were observed among the 50 haplotypes in the DIVS6 network (S3 Fig), approximately 39 unique haplotypes evolved from the common ancestors H5 and H1, reflecting distinct genetic makeup shaped by selective pressures. Eleven shared haplotypes descended from ancestral H2, indicating gene flow spreading susceptible and resistant alleles across disparate geographical regions. The haplotype network for the partial DIVS6 *vgsc* gene showed all haplotypes were clustered and closely related, regardless of carrying susceptible or resistant alleles.

#### Genetic diversity and neutrality test

The alignment of 90 individuals based on partial *vgsc* gene domains (DIIS6, DIIIS6, and DIVS6) revealed distinct genetic diversity indices and neutrality tests (Tajima’s D and Fu’s *Fs*) as tabulated in Table 2. Genetic diversity varied among geographical regions according to respective domains analysis. In Northern Peninsular Malaysia, genetic diversity indices revealed moderate nucleotide diversity (π) for DIIS6 and DIVS6, with values of 0.01201 and 0.01008, respectively, and a low π for DIIIS6 at 0.00823. The highest haplotype diversity (Hd) was observed in the DIVS6 analysis, with a value of 0.961. For demographic histories of *Ae. albopictus* populated in this region, neutrality test of DIIS6 indicated significant results, supporting the hypothesis of adaptive evolution in this partial gene region of *Ae. albopictus* (D = -1.92183; P < 0.01; Fs = -29.893). In contrast, the neutrality test results for DIIIS6 and DIVS6 insignificantly support the null hypothesis of adaptive evolution in these gene regions in this species. (DIIIS6: D = -0.94611; P<0.01 and *Fs* = -20.363, and DIVS6: D = -1.27244; P<0.01 and *Fs* = -33.646). These findings inferred the populations may have undergone recent expansion or experienced purifying selection [50,51].

**Table 2.**
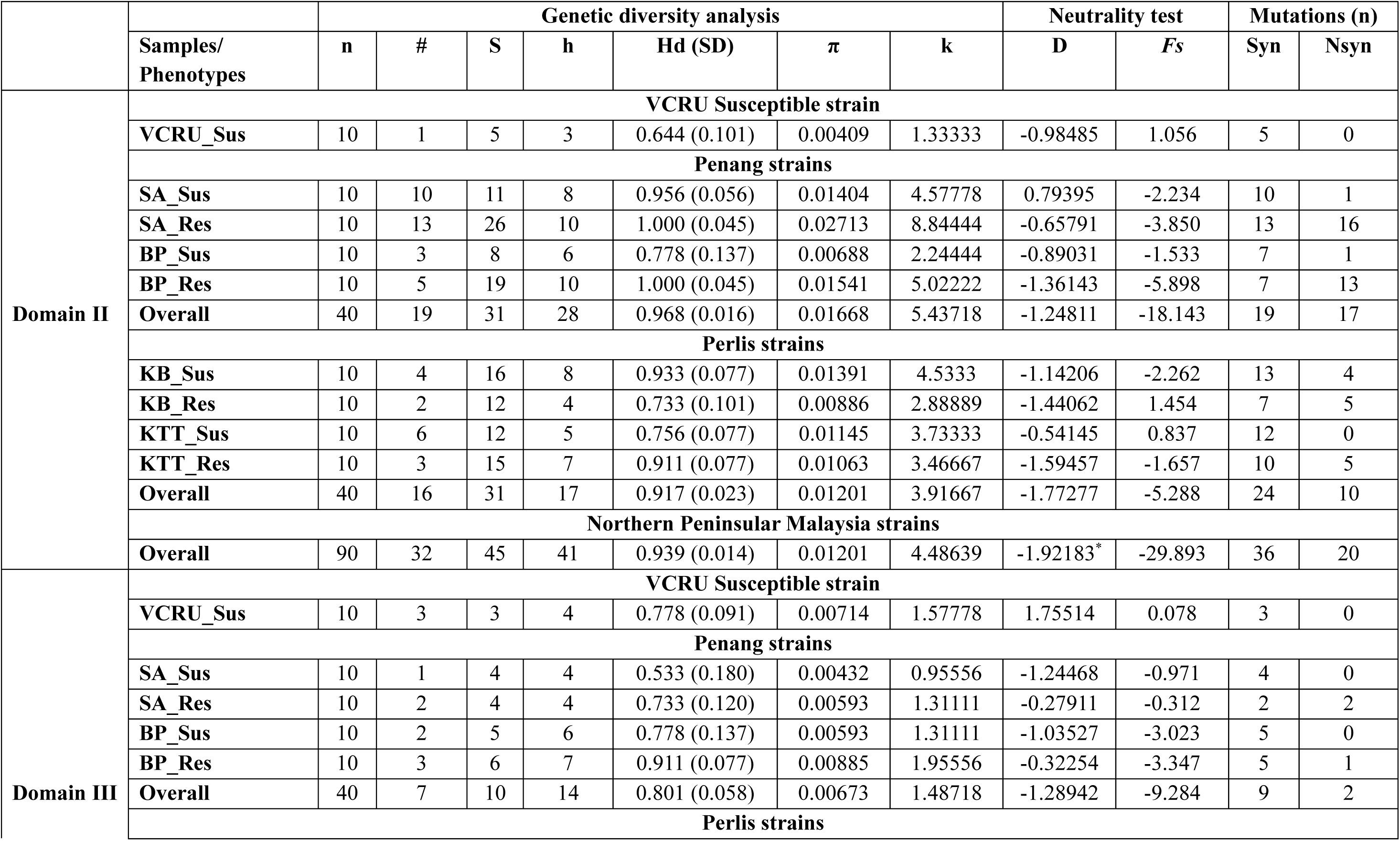
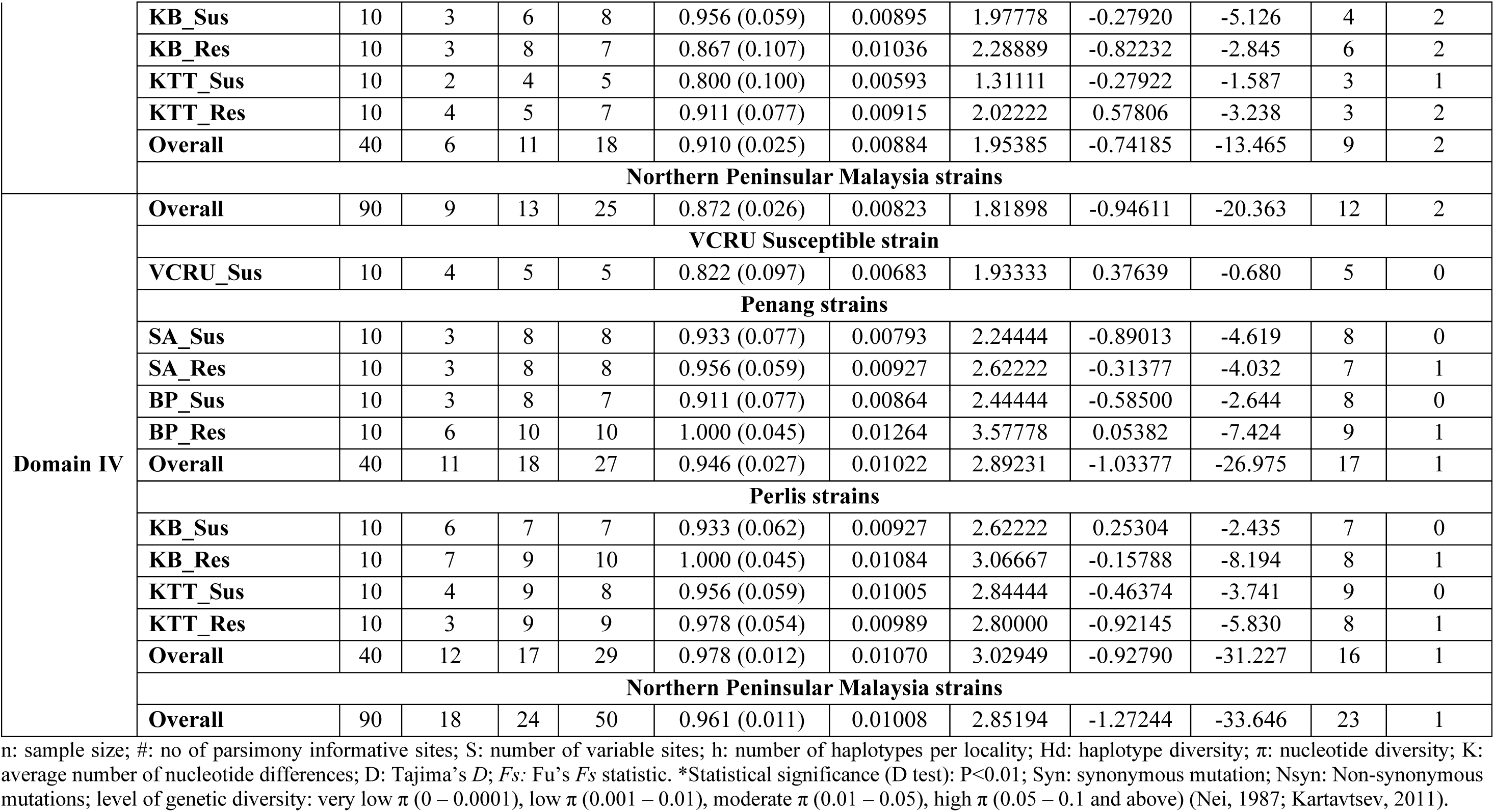
Genetic diversity, neutrality tests, and mutations in the partial *vgsc* gene domains between pyrethroids susceptible and resistant *Ae. albopictus* across Northern Peninsular Malaysia (Penang and Perlis).

### Protein prediction structures and docking of permethrin and deltamethrin to the partial DIIS6, DIIIS6, and DIVS6 *vgsc* gene of *Ae. albopictus*

#### Protein prediction structures

Homology modelling of the partial *vgsc* protein DIIS6, DIIIS6, DIVS6 were presented in wild-type, mutants, and combination alignments (wild-type versus mutant), denoting the mutation discovered in this study, namely A1022S/P, E1041K, F1534L, P1585K, and F1695L (S5 Fig). The interactions between the ligand (permethrin and deltamethrin) with the respective partial *vgsc* domain protein were analysed. Positional differences in ligand binding at the respective mutation sites are illustrated in S6 Fig, highlighting variations in binding positions compared to the wild-type.

#### Protein-ligand interactions

In-silico docking between the ligand and the studied *vgsc* protein revealed distinct affinity scores resulting from these interactions, as shown in Table 3. Protein prediction structures of DIIS6, DIIIS6, and DIVS6 with the lowest affinity interactions to permethrin and deltamethrin were portrayed in Fig 7. High affinity of insecticide to the *vgsc* binding site is determined by its low binding score value. The mutations in DIIS6 have caused an overall decreased in binding affinity for both ligands. Among these, mutant A (A1022S) exhibited the largest decrease in binding affinity, from -7.8 to -6.0 (kcal/mol) in the case of deltamethrin. Similarly, mutant B (A1022P), mutant C (E1041K), and the combined mutants A+C and B+C (A1022S + E1041K and A1022P + E1041K) also showed reduced binding affinities of -7.1 kcal/mol, -7.3 kcal/mol, -7.0 kcal/mol and -6.5 kcal/mol, respectively. In DIIIS6, mutant E (P1585R) exhibits slight increase in binding affinity, from -6.6 to -6.7 kcal/mol. On the other hand, mutant D (F1534L) and combined mutant D+E (F1534L + F1585R) both showed a decreased binding affinity of -6.2 kcal/mol.

**Fig 7.**
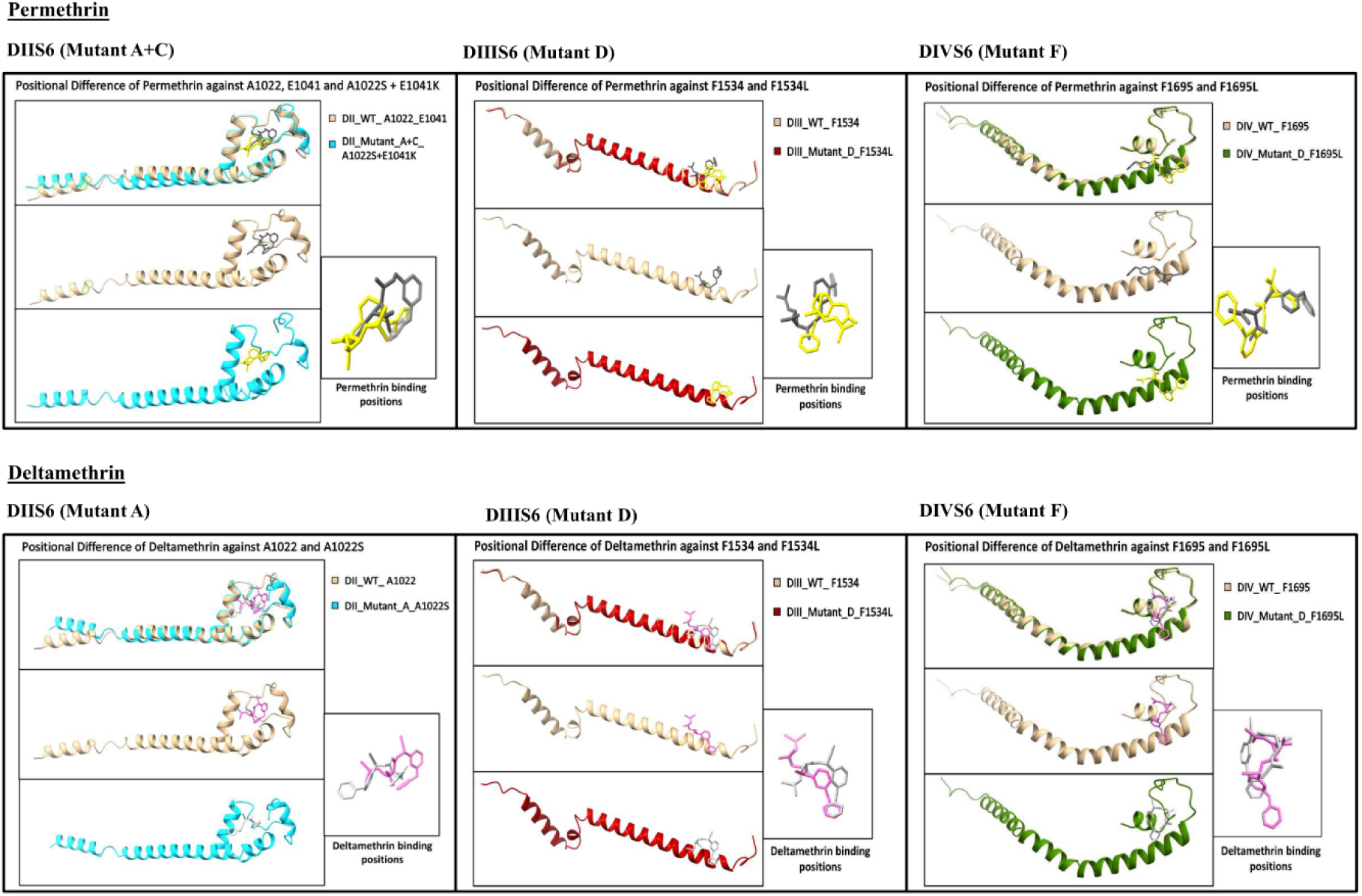
Protein-ligand interaction. Positional difference binding of permethrin and deltamethrin against the wild-type and mutant variants in DIIS6, DIIIS6, and DIVS6 with the lowest affinity interactions.

**Table 3.**
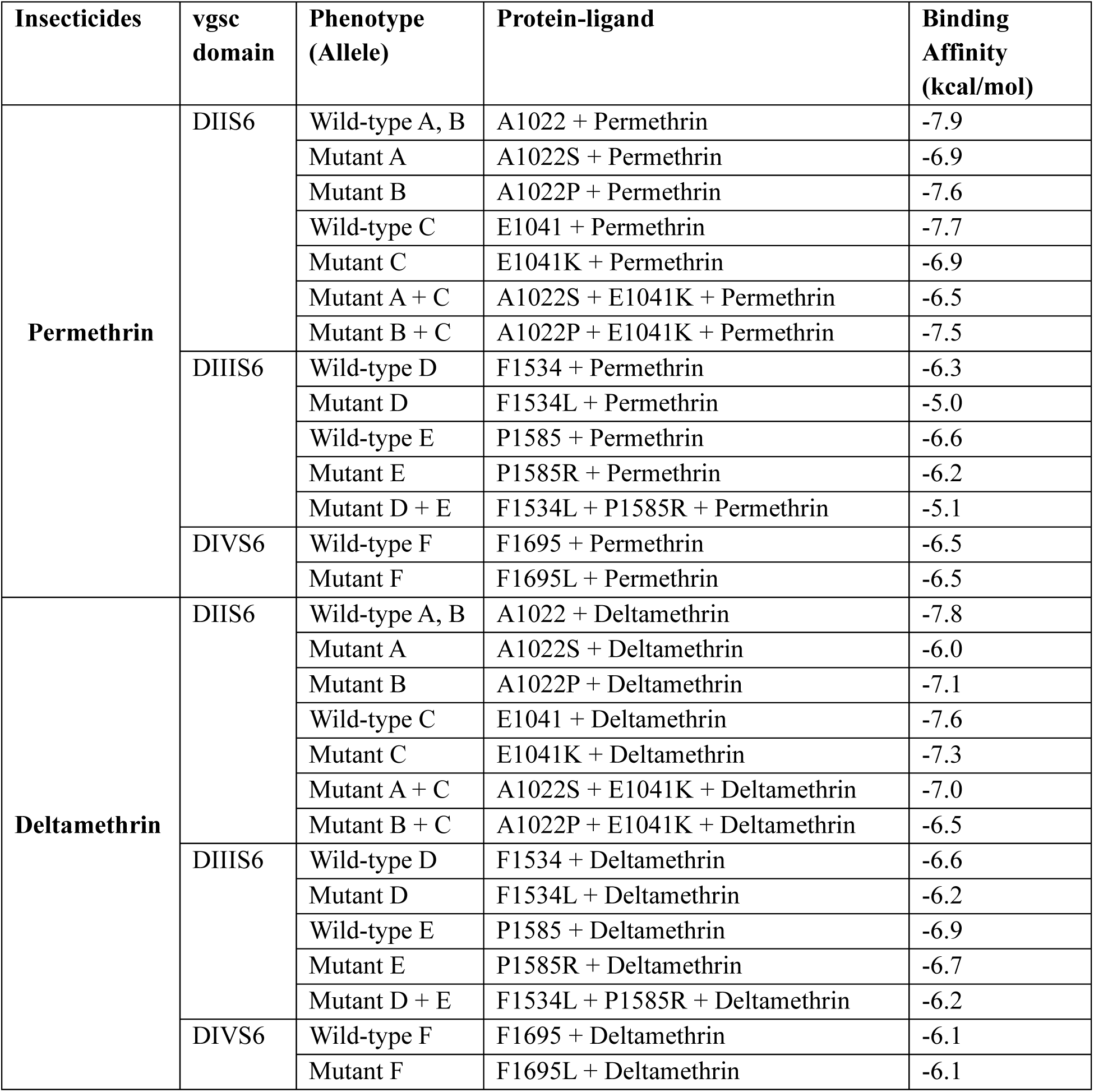
The binding affinity of insecticide to the A1022, E1041, F1534, P1585, and F1695 substituted *vgsc*. The binding energy was determined by AutoDock Vina scoring function.

For permethrin, mutations in DIIS6 also resulted in generally decrease in binding affinity, with mutant C (E1041K) and the combined mutants A+C (A1022S + E1041K) showing the most pronounced reduction, from -7.9 to -6.9 kcal/mol and -6.5 kcal/mol respectively. In contrast, mutant B (A1022P) and combined mutants B+C (A1022P + E1041K), display only slight reductions in binding affinity, to -7.6 and -7.5 kcal/mol respectively. In DIIIS6, mutant D (F1534L) and combined mutants D+E (F1534L + P1585R) caused significant drop in binding affinity, from -6.3 to -5.0 kcal/mol. The binding affinity in mutant E (P1585R) was slightly reduced to -6.2 kcal/mol. Finally, the F1695L mutation in DIVS6 exhibited no alterations in binding affinity for either deltamethrin or permethrin.

## Discussions

The chemical insecticide-based strategy has encountered obstacles over the years due to rising occurrence of insecticide resistance in mosquito vector populations [52]. The extent to which insecticide resistance influences the effectiveness of vector control programs remains a matter of contention in many countries nowadays. Likewise in Malaysia, fragmented and outdated data on insecticide resistance have raised significant concerns. Therefore, the present study was conducted to update data on insecticide susceptibility and to investigate the role of target-site mutations in conferring pyrethroid resistance in *Ae. albopictus* populations across different geographical settings in Northern Peninsular Malaysia.

### Resistance in Northern Peninsular Malaysian *Ae. albopictus*

The results underscore a major concern regarding the widespread resistance to permethrin, deltamethrin, primiphos methyl, and propoxur amongst adults, as well as resistance to temephos in *Ae. albopictus* larvae. In Malaysia, temephos is the primary larvicide used to treat mosquito larvae in stagnant water. Abate®, a potent temephos-derived larvicide, has been in use since 1973 [53], leading to the emergence of resistant larvae due to prolonged use of a single larvicide [54,55]. In addition, the widespread use of temephos larvicides in agricultural areas could reduce their effectiveness [56]. This is supported by the development of temephos resistance in KB and KTT strains of *Ae. albopictus* larvae, which thrive in areas with abundant stagnant water and rich microbiota, potentially impacting the larvicide’s residual effectiveness. Different levels of susceptibility to temephos may result from varied exposure histories [8], insensitivity of acetylcholine receptors (*Ace-1*) [57], and increased activity of metabolic detoxifying enzymes [58], all influenced by the genetic adaptation of mosquito species to selective pressures.

Adults *Ae. albopictus* from different regions in this study portrayed varying susceptibility patterns, regardless of dengue-prone classifications. This variability is due to the species utilized a diverse breeding sites and the different fitness costs from insecticide pressures within a population. The low mortality rates observed in the susceptibility bioassay for all four populations exposed to deltamethrin, permethrin, primiphos-methyl and propoxur suggest that they are resistant to these insecticides, as demonstrated in recent local studies [6,19,43]. This indicates that these insecticides are no longer effective in controlling these mosquito populations. The carbamate resistance pattern observed in this study is consistent with findings from Thailand, Malaysia, India, and China, where widespread propoxur resistance has been reported [16,43,59–63]. While carbamates are not the primary method for Malaysian dengue vector control, their extensive use in agriculture and public health over the past 50 years may have contributed to resistance [14,64]. Whilst the organophosphate insecticide used in this study serves as a substitute for malathion, which has been part of the vector control program since 2013, alternating with pyrethroids up to the present day [65]. The shift from malathion to the safer alternative, primiphos methyl, was due to resistance in *Aedes* mosquitoes. Therefore, the development of resistance to primiphos methyl, as seen in *Ae. albopictus* populations in this study, is not surprising.

Despite displaying resistance to pyrethroids, this study noted a contrast intensity of resistance towards permethrin and deltamethrin. Permethrin mediocrely striking down *Ae. albopictus* populations, with relatively moderate KdTs, assuring their ongoing use as a common residual spraying agent, especially in Penang. In contrast, deltamethrin was profoundly inefficient in affecting Perlis populations, amounting to the longest KdT_50_ and failing to achieve a high knockdown percentage (>5%). This indicates a significant degree of resistance, as deltamethrin could not produce a meaningful KdT_50_. Prolonged use of Type II pyrethroids like cypermethrin practiced by local health authorities as shared by Perlis Vector and Pest Unit in reducing the *Aedes* densities at most administrative zones in Perlis over years, justifying the current aftermath. Deltamethrin, sharing a cyano group with cypermethrin, interacts with the insect nervous system’s *vgsc* channel more strongly and persistently than permethrin, leading to higher toxicity [66]. Even so, this effectiveness can be altered by resistance mechanisms, as seen in certain populations. The higher mortality rates of KB and KTT strains to permethrin might suggest a reversion of their susceptibility upon contact with the older pyrethroid insecticides. Permethrin, conversely, has been extensively used in dengue vector control programs in Malaysia, with only a few instances of resistance reported nationwide [7,16,43,67]. Nonetheless, the establishments of pyrethroids-resistant *Ae. albopictus* in Perlis remarkably should not be overlooked, postulating the rotating different classes of insecticides are imperative. Our study depicted the heterogeneity in resistance degrees of *Ae. albopictus* from Penang, with emerging resistance to permethrin and deltamethrin, consistent with earlier findings by Chan et al. [68] and Zuharah et al. [69], albeit the differences in diagnostic doses. In contrast, Perlis strains displayed strong resistance to the same pyrethroid insecticides. Yet, there are no previous reports on susceptibility profiles of *Ae. albopictus* in Perlis prior to tracking their resistance histories. The differences of resistance pattern between Penang and Perlis based on permethrin and deltamethrin bioassays most possibly entail the different histories of pyrethroids spraying in these settings.

### Genotype-phenotype associations and their relationship to pyrethroid insecticides affinity

Resistance often involves the interaction of two or more mechanisms. In *Ae. albopictus*, pyrethroid resistance is frequently associated with both metabolic resistance and target-site resistance [70]. This species gained local attention after Ahmad et al. [19] identified the F1534L variant in permethrin-resistant *Ae. albopictus* (Kuala Lumpur strain), suggesting that sodium channel mutations indicate increasing target-site resistance in Malaysia. The F1534L mutation has been spreading in *Ae. albopictus* populations in Northern Peninsular Malaysia, with frequencies of 5% in Penang and 32.5% in Perlis, marking the second local report of this variant. Its origin in Malaysia remains unclear, requiring further investigation and broader screening across regions to better manage insecticide resistance. This mutation, identified as TTC to TTG or CTC, has also been reported globally over the past decade in locations such as Florida, USA (2011) [71], Italy (2011) [23], China (2014–2020) [24,31,34,64,72,73] and Indonesia (2024) [29]. This *kdr* mutation revealed a weak correlation (P<0.05) between heterozygous alleles (TTC/TTG) and resistance to permethrin and deltamethrin in Penang and Perlis populations, which aligns with other studies reporting no significant association between F1534L and resistance to these pyrethroids [31,72–74]. Conversely, other studies found that the F1534L mutation conferred protection against deltamethrin [62,75]. The weak correlation between heterozygous genotypes and resistance may result from mutant allele dosage, biochemical thresholds affecting susceptibility, and environmental pressures balancing resistance and fitness [76,77].

Multiple mutations A1022S/P, E1041K, P1585R, and F1695L, co-existed with F1534L in individual *Ae. albopictus* from Penang and Perlis (P<0.05) strongly influenced resistance to permethrin and deltamethrin, especially in dengue epidemic areas. This highlights the complex genetic interactions among non-synonymous mutations through epistasis driving insecticide resistance in these populations. Such results were reiterated in other studies, pointing out the possible epistatic effects contributing to resistance phenotypes in *Ae. albopictus* [29], and other species as well [78,79]. However, Modak and Saha [80] suggested that presence of other rare non-synonymous mutations in the *vgsc* gene of *Ae. albopictus* potentially compensate the fitness costs linked to *kdr* mutations, despite no observed fitness reduction. This could also explain the emergence of mutations such as A1022S/P, E1041K, P1585R, and F1695L, along with their distinct substitution patterns, in combination with the putative F1534 variant found in the studied species. Additional investigations into the genetic interactions among these mutations and their binding affinity scores are warranted.

Alignment and docking analysis of DIIS6 and DIIIS6 shows the reduction of binding affinity for permethrin and deltamethrin, even though the mutation sites were not located near the binding pocket. This suggests that the mutations affect ligand binding through allosteric effect. Permethrin and deltamethrin were likely to interact with the target protein through hydrophobic interactions. In the interactions of both ligands in DIIS6, the mutant A replaced alanine, a non-polar amino acid, with serine, which has a polar hydroxyl (-OH) group. This substitution introduces polarity into the protein, potentially disturbing the hydrophobic interactions towards both ligands. In mutant B, even though the replacement was made by a non-polar proline, the unique structure of proline that consist of a rigid cyclic structure may significantly disrupt the protein structure. This phenomenon was also described by Bajaj et al. [81] indicating that proline substitutions can alter protein structure, and was also shown in S6 Fig, showing significant structural changes compared to the wild-type protein. Whilst, in mutant C, the substitution of negatively charged glutamate with positively charged lysine reverses the charge, disrupting the overall protein charge and likewise interfering with ligand binding interactions.

The combined mutants A+C (A1022S + E1041K) in response to permethrin appear to combine the effects of altered polarity and charge reversal, further lowering the synergistic reduction in binding affinity from -6.9 to -6.5 kcal/mol. Mutants B+C (A1022P + E1041K) exhibited an additive effect, increasing the binding affinity from -7.6 kcal/mol in mutant B to -7.5 kcal/mol, although it remained lower than that of mutant C (-6.9 kcal/mol). Interestingly, deltamethrin elicited different responses in which mutant A+C exhibited additive-reduction effects, whereas mutant B+C displayed synergistic reduction. This likely stem from the chemical differences between the ligands.

Reviewing DIIIS6, mutant D substitutes phenylalanine with leucine at position 1534. Phenylalanine, an aromatic amino acid, is involved in hydrophobic stacking interactions [82]. The substitution of leucine and the absence of aromatic amino acid likely disrupt these interactions, leading to weaker binding affinity. Permethrin is slightly smaller and have lesser polar groups when compared to deltamethrin, thus may depend more on hydrophobic stacking. This may explain the larger reduction in affinity for permethrin (-6.3 to -5.0 kcal/mol) compared to deltamethrin (-6.6 to -6.2 kcal/mol), postulating this variant primarily affects the permethrin interaction. In mutant E, non-polar proline was replaced by positively charged arginine. Proline’s cyclic side chain added rigidity to the protein backbone, and its substitution with arginine may increase protein flexibility while altering its overall charge [81]. This could slightly disrupt the hydrophobic interactions with permethrin, leading to reduced binding affinity (-6.6 to -6.2 kcal/mol) and with deltamethrin (-6.9 to -6.7 kcal/mol). The combined mutants D+E (F1534L + P1585R) showed slight increase in the binding affinity for permethrin, from -5.0 kcal/mol for mutant D to -5.1 kcal/mol when combined with mutant E (-6.2 kcal/mol). Additionally, the combined mutants did not result in a further decrease in deltamethrin’s binding affinity. Such stability was likely due to the polar regions in deltamethrin compensating for the disruptions caused by the combined structural and charge changes introduced by both mutations. The substitution of phenylalanine with leucine at position 1695 (mutant F) in DIVS6 did not affect the binding affinity for either ligand. Despite phenylalanine’s role in hydrophobic stacking, the mutation’s distance from the binding site likely minimized its impact on interactions. The current findings provide the basic framework about the mutations and their interactions with pyrethroids; however, they lack comprehensive insights into these interactions across the full *vgsc* protein, emphasizing the need for further research.

### Impacts of insecticide-driven selection on the emergence of haplotypes and genetic variation

The extensive use of insecticides in agriculture for pest control and in vector management could exert significant selective pressure, driving the development of resistance in dengue vectors [83]. Insecticide resistance arises when the frequency of resistance genes in a mosquito population increases due to insecticide exposure, with selective pressure theoretically influencing the occurrence of mutations associated with resistance [52]. The discovery of *kdr* and rare synonymous mutations in the Northern Peninsular Malaysia regions postulates the effect of regional selective processes on the distribution patterns of existing haplotypes. Both Penang and Perlis populations share the same ancestral haplotypes, indicating that their wild-type genetic makeup was dominant before pyrethroid use, as shown in previous studies [23,84,85], regardless of the domains analyzed. The variations in haplotype networks based on the *vgsc* domains in this study may be attributed to differing evolutionary pressures acting on distinct regions of the gene. The *vgsc* gene comprises multiple domains, with mutations in specific regions significantly contribute to pyrethroid resistance under stronger selective pressure. Conversely, other regions experience less pressure, resulting in greater genetic diversity [86,87], in line with the currents findings.

Our findings suggest that haplotypes with the F1534L *kdr* allele, linked to resistance [61,75,29] can evolve independently in *Ae. albopictus* across diverse geographical regions, a common phenomenon in mosquitoes, including *Ae. albopictus* [23,85], *Ae. aegypti* [84,88], and *Anopheles sinensis* [89]. This independent evolution, potentially leading to adaptive evolution, highlights the emergence of new variant haplotypes in other domains, as discovered in this study. It raises uncertainty about whether the mutations originated independently in each region or spread between them, as identical mutations could suggest parallel evolution or gene flow. However, the unique *kdr* mutation patterns in resistant Perlis mosquitoes (H17 and H19) support the idea of localized, independent origins. P1585R haplotypes likely evolved from *kdr* haplotypes, enhancing resistance, mitigating fitness costs, or conferring cross-resistance to other insecticides. Their genetic divergence and reticulations suggest recombination events, forming new haplotypes, as observed in studies by Tancredi et al. [23], Zhang et al. [85], and Zhou et al. [89]. Similarly, this study found comparable evolutionary patterns in *Ae. albopictus*. Resistant *Ae. albopictus* haplotypes with the longest mutational edges evolved in both Penang and Perlis, indicating gene flow that allows resistance alleles to spread quickly. This rapid spread may be driven by migration, urbanization, and human activities that create interconnected breeding sites and adaptations [90], evoking recent research from neighbouring countries like Thailand. Given the recent expansion of *Ae. albopictus* beyond its native range, leading to the overlapping populations from different origins in new areas and the varied distribution of pyrethroid resistance mutations. On top of that, the regional distribution of *kdr* haplotypes emphasizes the need for ongoing monitoring of insecticide susceptibility to detect the emergence or spread of resistance in dengue vectors in Northern Peninsular Malaysia.

### Role of metabolic resistance in Northern Peninsular Malaysian *Ae. albopictus*

The effectiveness of PBO-synergists in counteracting pyrethroid resistance in *Ae. albopictus* from Penang suggests that metabolic resistance, involving cytochrome P450 monooxygenases and esterases, is prevalent in SA and BP strains. In Perlis populations, PBO-synergism showed varying success, with minimal restoration of susceptibility to deltamethrin compared to permethrin. This variation may result from differences in metabolic detoxification pathways, target-site mutations, or cuticular thickening. This observation is consistent with Koou et al. [91], who reported that the addition of PBO did not improve the effectiveness of cypermethrin and etofenprox against *Ae. aegypti* in Singapore. Similarly, PBO may be insufficient to restore susceptibility with deltamethrin, limiting its suitability for vector control in Perlis. Further biochemical and genome-wide transcription studies are demanded to clarify these findings. Nonetheless, the notable positive impact of PBO, particularly with permethrin, underscores its potential utility in managing insecticide resistance.

## Conclusion

Unravelling the updated susceptibility profiles of *Ae. albopictus* from Northern Peninsular Malaysia and the possible mechanisms of insecticide resistance revealed the species is resistant to permethrin, deltamethrin, primiphos-methyl, propoxur, and temephos. This study highlights the high prevalence of known mutations and identifies new point mutations in the *vgsc* gene of Malaysian pyrethroid-resistant strains. Our data supports the genetic exchange of *kdr* variant F1534L in other regions of Malaysia following its initial detection in Selangor/Kuala Lumpur. Regular surveillance of these mutations and their frequency is essential for detecting emerging resistance mechanisms. The add-in approaches such haplotype and in-silico docking studies shed light on distributions of pyrethroid-predictive mutations and their impact on pyrethroid target-site binding. Such comprehensive monitoring can guide effective insecticide use and resistance management, aiding local health authorities in planning more efficient vector control programs, particularly in Malaysia where epidemics can be highly destructive.

## Acknowledgments

The authors wish to thank local health authorities from Kangar District Health Office for helping with the mosquito sample collection. We thank Kenneth Tan JunKai for constructing the map for study location, and Ammar Khazaal Kadhim Almansoori for teaching and guiding on in-silico docking analysis.

## Author Contributions

**Conceptualization:** Nurul Adilah-Amrannudin.

**Sample collection:** Nurul Adilah-Amrannudin.

**Data curation:** Nurul Adilah-Amrannudin.

**Formal analysis:** Nurul Adilah-Amrannudin, Kenneth Tan JunKai.

**Funding acquisition:** Intan Haslina Ishak, Hasber Salim, Mohd Ghows Mohd Azzam.

**Investigation:** Nurul Adilah-Amrannudin.

**Methodology:** Nurul Adilah-Amrannudin, Kenneth Tan JunKai.

**Project administration:** Nurul Adilah-Amrannudin, Intan Haslina Ishak.

**Resources:** Nurul Adilah-Amrannudin, Intan Haslina Ishak, Hasber Salim, Nurul Nadia Manap, Shamsudin Abdul Rahman.

**Supervision:** Intan Haslina Ishak.

**Visualization:** Nurul Adilah-Amrannudin, Kenneth Tan JunKai.

**Writing – original draft:** Nurul Adilah-Amrannudin, Kenneth Tan JunKai.

**Writing – review & editing:** Nurul Adilah-Amrannudin, Intan Haslina Ishak, Hasber Salim, Abu Hassan Ahmad, Mohd Ghows Mohd Azzam.

## Supporting information

**S1 Table:** Lists of primers used in this study

**S2 Table: Mortality percentage of larval bioassay.** Percentage mortality of females *Ae. albopictus* at 24h post-treatment against diagnostic dose of 0.012 ppm and 0.034 ppm temephos larvicides, respectively.

**S3 Table: LC, DD, and RR of larval bioassay.** Summary of LC_50_, LC_95_, diagnostic dose (DD) and resistance ratios (RR) from larval bioassays on field strains of *Ae. albopictus* against USM VCRU laboratory strain.

**S4 Table: Mortality percentage of adult bioassay.** Percentage mortality of females *Ae. albopictus* at 24h post-treatment against several insecticides from carbamate, organophosphate and pyrethroid classes, together with 4% PBO as a synergist for pyrethroid insecticides.

**S5 Table: Knockdown rates.** Knockdown rates of *Ae. albopictus* in Penang and Perlis, Malaysia against pyrethroid adulticides.

**S6 Table: Genotypic distributions.** Distributions of prevalent mutations in *vgsc* gene domains of pyrethroids susceptible and resistant *Ae. albopictus* across Northern Peninsular Malaysia (NPM) populations.

**S7 Table: Genotype-phenotype association.** Association of genotype F1534L with the resistance phenotype towards pyrethroid insecticides for the Penang and Perlis strains of *Ae. albopictus*.

**S8 Table: Genotype-phenotype association.** Association of genotype A1022S/P with the resistance phenotype towards pyrethroid insecticides for the Penang and Perlis strains of *Ae. albopictus*.

**S9 Table: Genotype-phenotype association.** Association of genotype E1041K with the resistance phenotype towards pyrethroid insecticides for the Penang and Perlis strains of *Ae. albopictus*.

**S10 Table: Genotype-phenotype association.** Association of genotype P1585R with the resistance phenotype towards pyrethroid insecticides for the Penang and Perlis strains of *Ae. albopictus*.

**S11 Table: Genotype-phenotype association.** Association of genotype F1695L with the resistance phenotype towards pyrethroid insecticides for the Penang and Perlis strains of *Ae. albopictus*.

**S12 Table: Genotypic combinations.** Distribution of single-, duo-, triple-, and quadruple-loci of the genotypic combination in DIIS6, DIIIS6, and DIVS6 of *vgsc* gene in *Ae. albopictus*.

**S13 Table: Haplotypes frequency.** A) Forty-one haplotypes were identified from 90 nucleotide sequences of DIIS6 in *vgsc* gene fragment of local *Ae. albopictus* in Penang and Perlis, and deposited in the NCBI Genbank. B) Twenty-five haplotypes were identified from 90 nucleotide sequences of DIIIS6 in *vgsc* gene fragment of local *Ae. albopictus* in Penang and Perlis, and deposited in the NCBI Genbank. C) Fifty haplotypes were identified from 90 nucleotide sequences of DIVS6 in *vgsc* gene fragment of local *Ae. albopictus* in Penang and Perlis, and deposited in the NCBI Genbank.

**S1 Fig: Ramachandran plot.** Structural validation based on Ramachandran plot analysis.

**S2 Fig: Graph of mortality rate of *Ae. albopictus* larvae.** Percentage mortality of *Ae. albopictus* larvae from the different localities in Penang and Perlis, Malaysia towards 0.012 ppm and 0.034 ppm of temephos. Error bars represent 95% confidence interval (CI). Black horizontal line indicates resistant threshold level (mortality <90% is considered phenotypically resistant; WHO, 2016).

**S3 Fig: Haplotype network of DIIS6.** Haplotype network showing the genealogical relationships of 41 haplotypes in DIIS6 based on four strains of Northern Peninsular Malaysia populations of *Ae. albopictus*: wild-type haplotypes are depicted in black and those with new non-synonymous mutations are highlighted in blue.

**S4 Fig: Haplotype network of DIVS6.** Haplotype network showing the genealogical relationships of 50 haplotypes in DIVS6 based on four strains of Northern Peninsular Malaysia populations of *Ae. albopictus*: wild-type haplotypes are depicted in black and those with new non-synonymous mutations are highlighted in blue.

**S5 Fig: Structural alignment of partial *vgsc* protein domains.** A) Structural alignment of partial *vgsc* gene domains DIIS6 between wild-type and mutant variants. B) Structural alignment of partial *vgsc* gene domains DIIIS6 between wild-type and mutant variants. C) Structural alignment of partial *vgsc* gene domains DIVS6 between wild-type and mutant variants.

**S6 Fig: Protein-ligand interactions.** A) Positional difference binding of permethrin against the wild-type and mutant variants in DIIS6. B) Positional difference binding of permethrin against the wild-type and mutant variants in DIIIS6. C) Positional difference binding of permethrin against the wild-type and mutant variants in DIVS6. D) Positional difference binding of deltamethrin against the wild-type and mutant variants in DIIS6. E) Positional difference binding of deltamethrin against the wild-type and mutant variants in DIIIS6. F) Positional difference binding of deltamethrin against the wild-type and mutant variants in DIVS6.

